# Predicting developmental relationships of tumor resident and circulating T cells in ovarian cancer

**DOI:** 10.1101/2023.02.28.530488

**Authors:** Mayra S. Carneiro, Yacine Bareche, Cheng Zhao, Pamela Thébault, Kurosh Rahimi, Diane Provencher, Vanessa Samouélian, Béatrice Cormier, Jean-François Cailhier, Anne-Marie Mes-Masson, Sophie Petropoulos, John Stagg, Réjean Lapointe

## Abstract

Characterizing T cell populations and understanding their developmental relationships may help design more effective cancer immunotherapies. We coupled single-cell transcriptomics and T cell receptor (TCR) αβ profiling of intratumoral and peripheral T cells in ovarian cancer patients to identify transcriptional programs and infer their relationship by trajectory and TCR overlap analyses. We proposed a model of differentiation pathway from an intermediate GZMH-expressing CD8 T cell subset found in the blood and tumor that progressively reinforces the exhaustion and tissue residency programs from a *CCL4*-expressing cluster towards *XCL1*- and *CXCL13*-expressing terminally exhausted cells. Inferred cell communication analysis suggests that interaction with *CXCL13*-expressing CD4 T cells, which we refer to as Tfh-like cells, sustains the effector function of this intermediate GZMH-expressing CD8 T cell subset. Moreover, our results suggest that Tfh-like cells attract cells expressing *GPR183* through the production of its ligand 7α,25 dihydroxycholesterol (7α,25-HC). Finally, we demonstrated that *GPR183* is highly expressed in a subset of pre-effector *GZMK*-expressing CD8 T cells and plasmacytoid dendritic cells. Collectively, our results suggest that Tfh-like cells expressing IL-21 help promote antitumor immunity against ovarian tumors by coordinating the action of immune cells responsive to 7α,25-HC.

## INTRODUCTION

Although tumor infiltration with lymphoid cells is associated with a better prognosis in several cancer types, immunotherapy targeting immune checkpoint receptors only benefits a subgroup of patients. Deep profiling of CD8 T cell subpopulations responsive to immune checkpoint blockade (ICB) may help us better understand the mechanisms of resistance and improve treatment efficacy. It is well established that chronic infections and cancer promote CD8 T cell exhaustion characterized by the loss of effector function, high expression of immune-regulatory checkpoint receptors, as well as epigenetic and metabolic reprogramming (*1*). During this process, the transcriptional factor TCF-1 maintains self-renewal (i.e., stemness) of precursor-exhausted T cells (T_PEX_), while the transcriptional factor TOX commits T_PEX_ to the exhaustion program (*2*–*7*). Interestingly, T_PEX_ are present in several cancer types (*8*) and predict ICB response (*9*). Antitumor responses from T_PEX_ rely on their ability to undergo self-renewal and differentiate into effector cells (*10, 11*). Conversely, terminally differentiated exhausted CD8 T cells (T_TERM_) show increased levels of inhibitory receptors, diminished effector function, and acquire transcriptional programs involved in tissue retention (*11, 12*). Developmental relationships of four subsets of exhausted CD8 T cells have been recently demonstrated in a mouse model, but understanding their transitional states in human tumors remains challenging (*11*).

Initially, it was proposed that ICB could reinvigorate intratumoral T_TERM_ cells, but recent studies on T cell dynamics suggest that antitumor immune responses rely on replacing tumor-resident T cells with peripheral T cells (*13, 14*). Moreover, the blockade of PD-1 in the tumor-draining lymph nodes (TDLN), rather than the tumor site, has been shown to promote activation and proliferation of T_PEX_ cells, which then migrate to the tumor (*15, 16*) and settle in regions enriched with antigen-presenting cells, reminiscent of T cell zones of secondary lymphoid organs (SLO) (*10*).

The presence of lymphoid aggregate, called tertiary-lymphoid structures (TLS), is associated with better ICB responses (*17*–*20*) and is believed to be the site for local antibody production (*21, 22*) and T cell priming (*23*). TLSs arise in non-lymphoid tissues and resemble lymph follicles in the SLO with segregated T and B cell zones. Although molecular mechanisms of TLS formation are similar to SLO, the cellular drivers are partly different. For instance, the chemokines required in TLS formation (such as CCL19, CCL21, and CXCL13) are produced by local cells instead of lymphoid tissue inducer (LTi) and lymphoid tissue organizer (LTo) cells observed in lymphoid tissue organogenesis (*24, 25*). Central to TLS formation are T follicular helper cells (Tfh), which mediate the recruitment of B cells by producing CXCL13 and enhance the cytotoxic function of CD8 T cells at the site of infection (*26, 27*) and tumors (*28*–*32*).

Here we use scRNA-seq to deeply characterize intratumoral and peripheral CD45+ immune cells in patients with high-grade serous ovarian carcinoma (HGSOC), the most aggressive and lethal subtype of ovarian cancer (*33*). By studying the transcriptome, TCR repertoire, and location of tumor-infiltrating and circulating T cells, we tracked T cell clones expanded in the periphery and tumor and deciphered changes in cell state triggered by entering the tumor tissue. We identified a subset of tumor-infiltrating CD4^+^ T cells that upregulate *CXCL13*, which we refer to as T follicular helper-like (Tfh-like) cells, that associate with increased lymphoid cell recruitment to ovarian tumors. Our data strongly suggest that Tfh-like cells participate in the recruitment of CXCR5-expressing B cells, produce IL-21 to support the effector function of dual-expanded CD8 T cells, and process cholesterol into 7a,25-dihydroxycholesterol (7a,25-HC) involved in the recruitment of GPR183-expressing cells and coordination of B cell, T cell, and plasmacytoid dendritic cells in lymphoid aggregates.

## RESULTS

### T cell heterogeneity

We used scRNA-seq to simultaneously study transcriptome and TCR-seq of CD45+ tumor-infiltrating immune cells from five patients with HGSOC and matched blood-derived immune cells from four of these patients. To understand the heterogeneity of T cells, we investigated their gene expression profiles, inferred location, and diversity of TCRαβ repertoire. PBMCs from two age-matched healthy donors served as controls. We independently analyzed immune cell populations in blood and tumors (fig. S1A) and selected the cells within clusters annotated as T cell (clusters expressing the genes *CD3G, CD3D, CD8A*, and *CD4*) to further integrate the 47,182 cells into a separate unsupervised clustering analysis (Fig. 1A). To confirm T cell annotation, we looked at the level of expression of CD3G and the frequency of TCRαβ sequenced in each cluster (fig. S1B).T cells from tumor and blood were distributed across 20 clusters (Fig. 1B), of which six were CD4 cells (CD4_CCR7, CD4_KLRB1, CD4_NEAT1, CD4_FOS, CD4_ISG15, and CD4_FOXP3), eight were CD8 cells (CD8_GZMB, CD8_GNLY, CD8_GZMK, CD8_CCL4, CD8_GZMB, CD8_ZNF683, CD8_XCL1, and CD8_CXCL13) and two other clusters defined as under stress (i.e. ‘HS’; heat-shock) or mitosis (i.e. ‘prolif’). Both tumor and blood were enriched for CD4 T cells; in blood, the CD4_CCR7 cluster was the most abundant, while in tumors, the CD4_FOXP3 cluster was most abundant (fig. S1C). To characterize T cell states, we performed marker detection (table S2 and fig. S1D) and named clusters based on one of the most expressed genes (Fig. 1C and D). Using this approach, we identified the CD8_GZMB and CD8_GZMH clusters are highly expressing multiple genes associated with T cell effector function (e.g., *GZMA, GZMH*, and *PRF1*), and the CD8_XCL1 and CD8_CXCL13 clusters expressing genes related to terminal exhaustion (e.g., *PDCD1, HAVCR2, TOX*, and *CXCL13*).

**Fig. 1.**
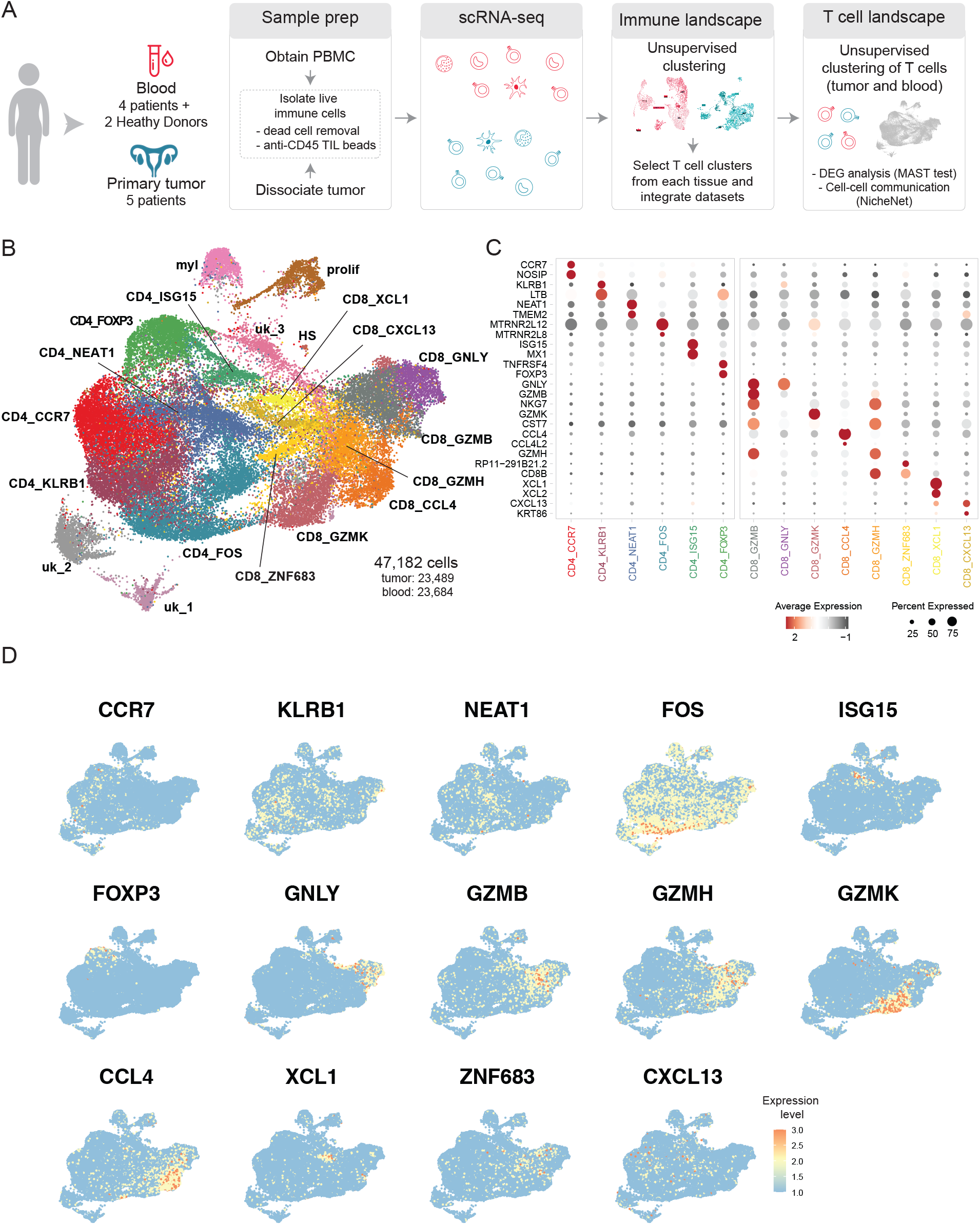
T cell landscape in high-grade serous ovarian carcinoma (HGSOC). **(A**) Schematic experiment design including samples from five untreated high-grade serous ovarian cancer patients and two healthy donors. Blood and tumor samples were prepared and analyzed as described in the diagram. The T cells identified in each tissue were selected and integrated to obtain the T cell landscape. (**B**) Unsupervised clustering analysis on 47,182 T cells (23, 489 intratumoral and 23,684 peripheral) represented in Uniform Manifold Approximation and Projection (UMAP). (**C**) Dot plot of the top two markers identified in the differential expression genes analysis by comparing each cluster to all other cells. (**D**) UMAP projection showing the expression level of genes used to name the cluster.

### A core signature of T cell exhaustion defines TILs

To identify cell states associated with the tumor microenvironment, we evaluated the proportion of cells from blood or tumor and the abundance of clusters within those tissues. Cells from tumors were mainly distributed across clusters CD8_CXCL13, CD8_XCL1, and CD8_CCL4 (Fig. 2A), and tumors were enriched with CD4_FOXP3 cells (fig. S1C). Genes related to exhaustion (*TOX, PDCD1, ENTPD1, CXCL13, HAVCR2*) were the core transcriptional signature of intratumoral CD8 T cells (Fig. 2B-C). Notably, these genes are known to be highly expressed in terminally-exhausted T cells across several cancer types (*12, 34*–*37*), and were highly expressed in our CD8_CXCL13 and CD8_XCL1 clusters (Fig. 2B).

**Fig. 2.**
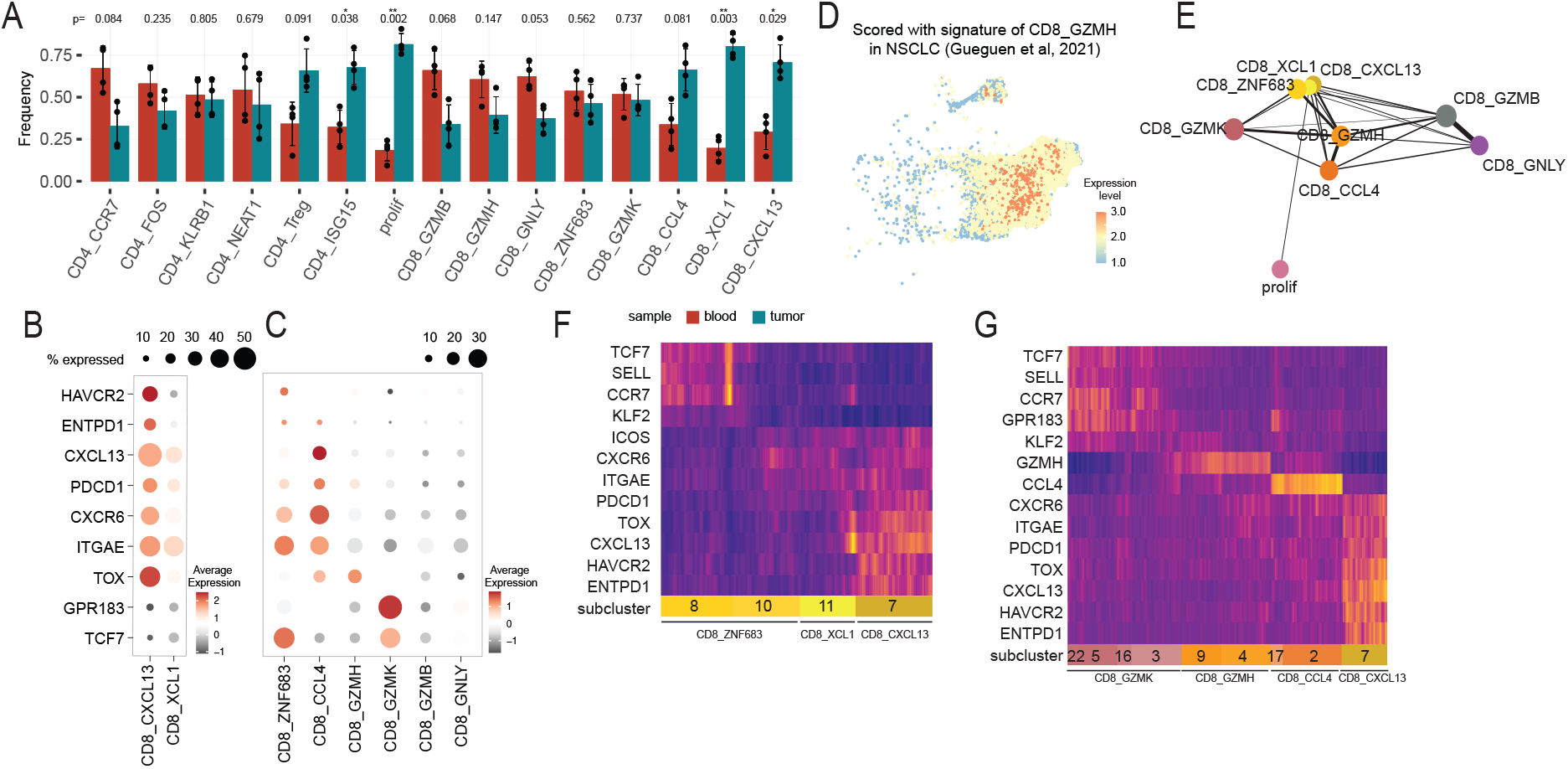
States of tissue-resident exhausted CD8 T cells and memory-like cells in HGSOC. (**A**) Bar plots showing the mean frequency of cells from blood (red) or tumor (blue) in each cluster. Data are expressed as mean ± SD of individual patient. Significant p values determined by the t statistics. (**B-C**) Dot plots of normalized average expression of genes related to exhaustion (*TOX, PDCD1, ENTPD1, CXCL13*, and *HAVCR2*), tissue residence (*ITGAE* and *CXCR6*), and memory (*TCF7*), including all CD8 T cell clusters (**B**) and excluding clusters CD8_CXCL13 and CD8_XCL1 (**C**). (**D**) UMAP projection of CD8 T cells scored with the signature of transitional CD8 T cell cluster identified in lung cancers (*37*). The color scale represents low (blue) to high (red) scores. (**E**) Trajectory analysis using partition-based graph abstraction (PAGA) on CD8 T clusters. (**F and G**) Heat maps of gene expression level along the PAGA paths in CD8_ZNF683 (**F**) and CD8_GZMK (**G**).

Interestingly, the CD8_GZMH cluster showed higher expression of genes related to cytotoxicity (*GZMH, GZMB, NKG7*) and early stage of exhaustion (*TOX* and *PDCD1*), suggesting a transitional state between effector and exhaustion. Of note, a previously reported transitional signature in NSCLC highly scored with cells in the CD8_GZMH cluster in our dataset (Fig. 2D). Therefore, we hypothesized that ovarian tumors harbor transitional states of exhausted T CD8 cells. To test this hypothesis, we inferred cell trajectory using partition-based graph abstraction (PAGA) (*38*). PAGA analysis projected CD8 clusters in six branches (Fig. 2E), where CD8_GZMH was settled in the center, connecting with all other clusters, especially with CD8_CCL4, CD8_ZNF683, CD8_GZMK, CD8_CXCL13, and CD8_GZMB clusters (the weighted edges represent the connectivity between clusters). Together, these results suggest CD8_GZMH is a transitional state between effector and exhausted cells, with a gradual increase in expression of genes related to exhaustion and tissue-residency in CD8_ZNF683, CD8_CCL4, CD8_XCL1, and CD8_CXCL13.

In addition to localization and gene expression profile, the T cell repertoire may also help us understand the states and dynamics of T cells in HGSOC patients. Therefore, we investigated the diversity, expansion, and clonal overlap of T cell clusters. Heterogeneity calculated by D50 index score the least diversity cluster close to zero, corresponding to CD8_GZMB, CD8_GZMH, and CD8_CXCL13 (Fig. 3A). Clones expanded in both tissues (dual-expanded) were enriched in effector-like clusters (CD8_GZMB and CD8_GZMH), whereas tumor-expanded clones are more frequent within CD8_CXCL13 and CD8_XCL1 (Fig 3C-D). Moreover, we proposed the CD8_CCL4 cluster as an intermediary between dual-expanded and tumor-expanded states once clones classified in these groups are equally frequent and the TCR overlap in extension greater than any other cluster (Fig. 3B). CD8_ZNF683 has a similar proportion of tumor-expanded clones; however, dual-expanded clones were poorly present. Together, gene expression, location, and TCR repertoire of T cells in our dataset suggest a gradual acquisition of exhaustion program initiated by upregulation of *TOX* and *PDCD1* in effector-like cells (e.g., CD8_GZMH). This cluster was represented by a large expansion of dominant clones from blood and tumor origin. Following, the CD8_CCL4 cluster increased the proportion of tumor-expanded clones and the expression of genes related to exhaustion (e.g., *ENTPD1, HAVCR2*, and *CXCL13*) and tissue retention (e.g., *CXCR6* and *ITGAE*) (Fig. 2C). In the subsequent states (CD8_XCL1 and CD8_CXCL13), the level of expression of those genes and the proportion of tumor-expanded clones increased, while the TCR diversity decreased, suggesting that few clones progressed to terminal exhaustion state. These results suggest a hierarchical developmental pathway for exhausted CD8 T cells initiated in CD8_GZMH, followed by CD8_CCL4, CD8_XCL1, and CD8_CXCL13 clusters.

**Fig. 3.**
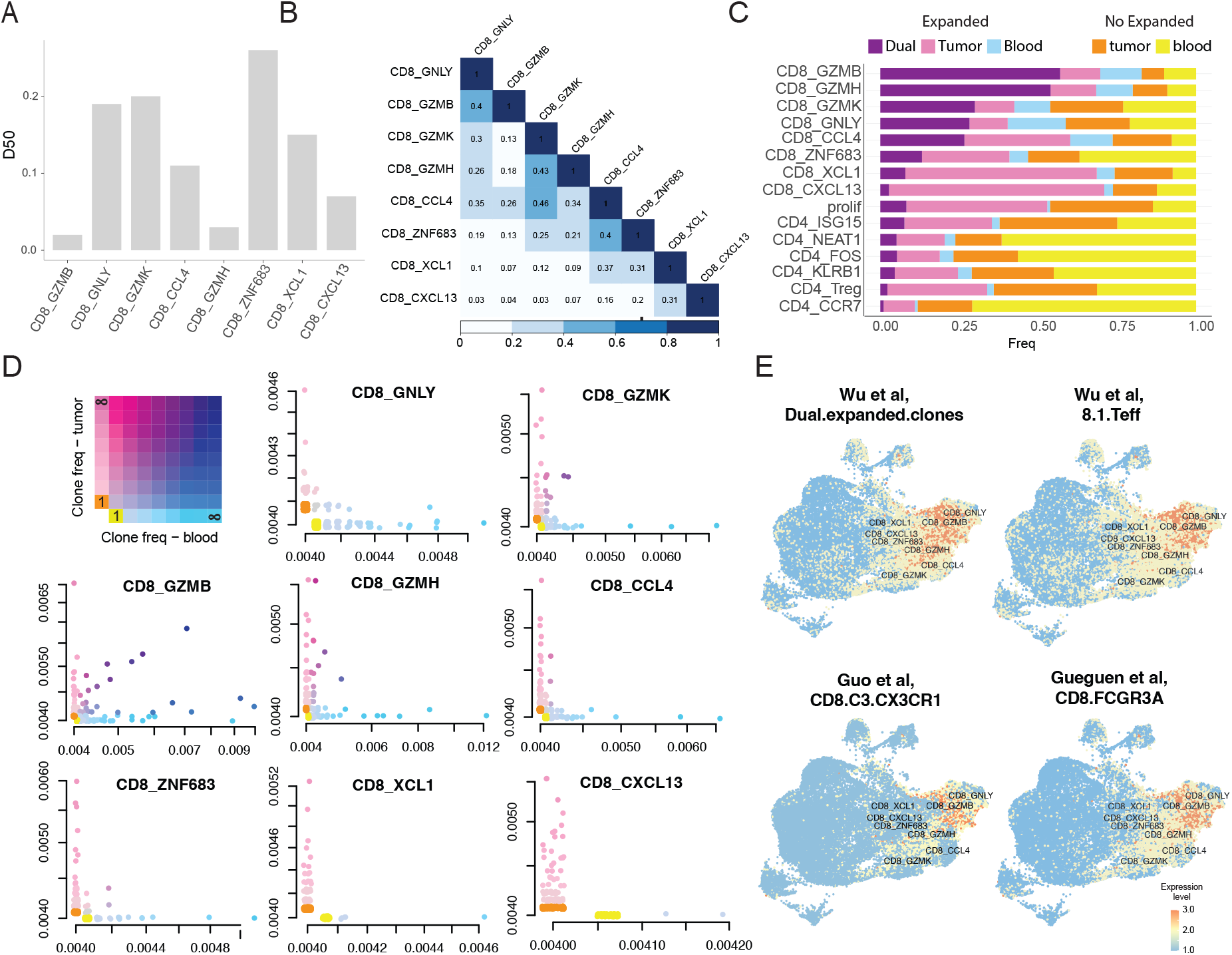
Dual-expanded clones are effector-like CD8 T cells. To investigate the spatial expansion of TCRs, we classified the clones of T cells into five groups as proposed by Wu et al.: singleton within the blood (orange) or tumor (yellow), multiplets restricted to blood (light blue) or tumor (pink), or expanded in both tissues (gradient between pink and blue). (**A**) The bar plot of the diversity index shows the percentage of dominant clones that account for 50% of each cluster’s total number of CD3Rs. (**B**) Heat map representing the TCR overlap calculated using the Morisita-Horn index. (**C**) Bar plot indicating the proportion of five types of clones classified by the spatial expansion criteria in each cluster. (**D**) The first plot is a schematic view of scatter plots representing the frequency of clones classified at the cluster level instead of the patient’s sample level shown in C. For each cluster, we showed the number of dual-expanded clones (Nd) and Pearson’s correlation coefficient measuring the relationship between the frequency of a dual-expanded clone in the blood and tumor. (**E**) UMAP projection of T cells scored with external signatures of effector-like cells or dual-expanded clones (*14, 34, 37*). The color scale represents low (blue) to high (red) scores.

### Two subsets of stem-like memory CD8 T cells in HGSOC

It has been proposed that therapeutic responses to ICB rely on the expansion of stem-like memory CD8 T cells, also known as precursor-exhausted cells (T_PEX_). This population expresses the transcriptional factor TCF-1 and has the ability to self-renew and differentiate into effector cells. We thus evaluated the expression of *TCF7* (gene encoding TCF-1) in our dataset and identified potential T_PEX_ within clusters CD8_GZMK and CD8_ZNF683 (Fig. 2C). To better characterize these cells, we defined subclusters in PAGA projection. This revealed two subsets of memory-like cells within ovarian cancer patients: k_22 (from CD8_GZMK) and k_8 (from CD8_ZNF683) (fig. S2A). These clusters expressed high levels of genes associated with circulating memory cells, including *SELL* and *CCR7*, and, intriguingly, cluster k_22 expressed a high level of exhaustion-related genes (*CTLA4, TIGIT*, and *HAVCR2*) as well as *GPR183* (fig. S2B). GPR183 encodes for EBI2, the receptor for the oxysterol 7α, 25-OHC, and is involved in the positioning of B cells in follicular regions of the lymph node (*39*).

We next characterized the k_22 and k_8 subclusters regarding the tissue of origin, the gene expression, and the TCR repertoire. Our analysis suggests that the two subclusters represent cells originating from the blood (fig. S2C) that diverged into tissue-resident or granzyme K lineages. Accordingly, the trajectory analysis and gene signature (Fig 2F, fig. S2A, and table S3) showed k_8 in the tissue-resident program with gradual upregulation of *CXCR6* and *ITGAE*, and downregulation of *KLF2* following the trajectory towards k_10, another subcluster from CD8_ZNF683. This lineage increased the expression of inhibitory receptors and genes associated with exhaustion as it moved toward subclusters k_11 (cluster CD8_XCL1) and k_7 (cluster CD8_CXCL13). The TCR repertoire confirmed this trajectory as clones of k_8 mostly overlapped with k_10, k_11, and k_10 (fig. S2D). It also shared clones with k_2, a subcluster of CD8_CCL4, suggesting plasticity between terminally-exhausted CD8 T cells and CD8_CCL4 phenotype. On the other hand, k_22 appears to differentiate toward CD8_GZMK, including subclusters k_5, k_16, and k_3. The trajectory analysis and the gene expression profile showed that as they moved toward k_3, they upregulated markers of effector-like cells subclusters (k_14 from CD8_CCL4, and k_9 from CD8_GZMH), such as *GZMM, CD8A*, and *CST7* (Fig. 2F and fig. S2B). Importantly, this subset of CD8 T cells expressing high levels of GZMK has been recently described as pre-effector T cells (*40*), supporting the differentiation pathways suggested in our analysis. Therefore, we appear to have identified two subsets of precursor memory-like CD8 T cells within ovarian cancer patients, one within pre-effector T cells that upregulate exhaustion-related genes and another from CD8_ZNF683 committed to the tissue-resident program.

### Dual-expanded clones interact with T Follicular helper-like (Tfh-like) cells in the TME

The presence of T cell clones expanded in both tumor and peripheral tissue, also called dual-expanded T cells, can predict response to ICB (*14*). Dual-expanded clones are more frequent in CD8_GZMB and CD8_GZMH clusters. To confirm this association at the gene expression level, we evaluated the expression of the gene signature of dual-expanded clones from Wu et al. in our dataset (Fig. 3E). The cells from clusters CD8_GZMB and CD8_GZMH indeed highly scored with the gene signature of dual-expanded clones. We further compared our dataset with signatures of effector-like cells from other studies (*34, 37*), and again observed an association (Fig. 3E).

Considering the importance of dual-expanded clones in antitumor immune response, we next interrogated the cellular interactions they encountered in the tumor microenvironment. We choose the tool NicheNet (*41*) because instead of restricting the analysis to ligand and receptor expression, as most of the other tools, NicheNet considers the signal transduction and target genes to predict receptors modulating the phenotype. Therefore, in our analysis, we defined sender cells as those from the intratumoral immune cells dataset (fig. S3A) and the receiver cells as those dual-expanded in CD8_GZMB or CD8_GZMH clusters. Using this approach, we identified the ligands present in the tumor-immune microenvironment (TIME) modulating transcriptional changes in dual-expanded clones from CD8_GZMB (fig. S3B) or CD8_GZMH (Fig. 4A and fig. S3C). One of the top ligands in both analyses was *IL21*, the gene encoding for a cytokine known to sustain the effector function of CD8 T cells (*23, 30*). We accessed the source of *IL21* in the transcriptome data of TIME (Fig. 4A), which identified a subset of CD4 cells (cluster CD4_CXCL13) (Fig. 4B). Interestingly, those cells were already described in non-lymphoid tissues as Tfh-like cells, CXCL13-producing CD4 T cells, or T resident helper cells (*26, 27, 32, 42*). Thus, our interaction analysis suggested that *IL21* expressed by CD4_CXCL13 modulates effector-like dual-expanded clones in the TME.

**Fig. 4.**
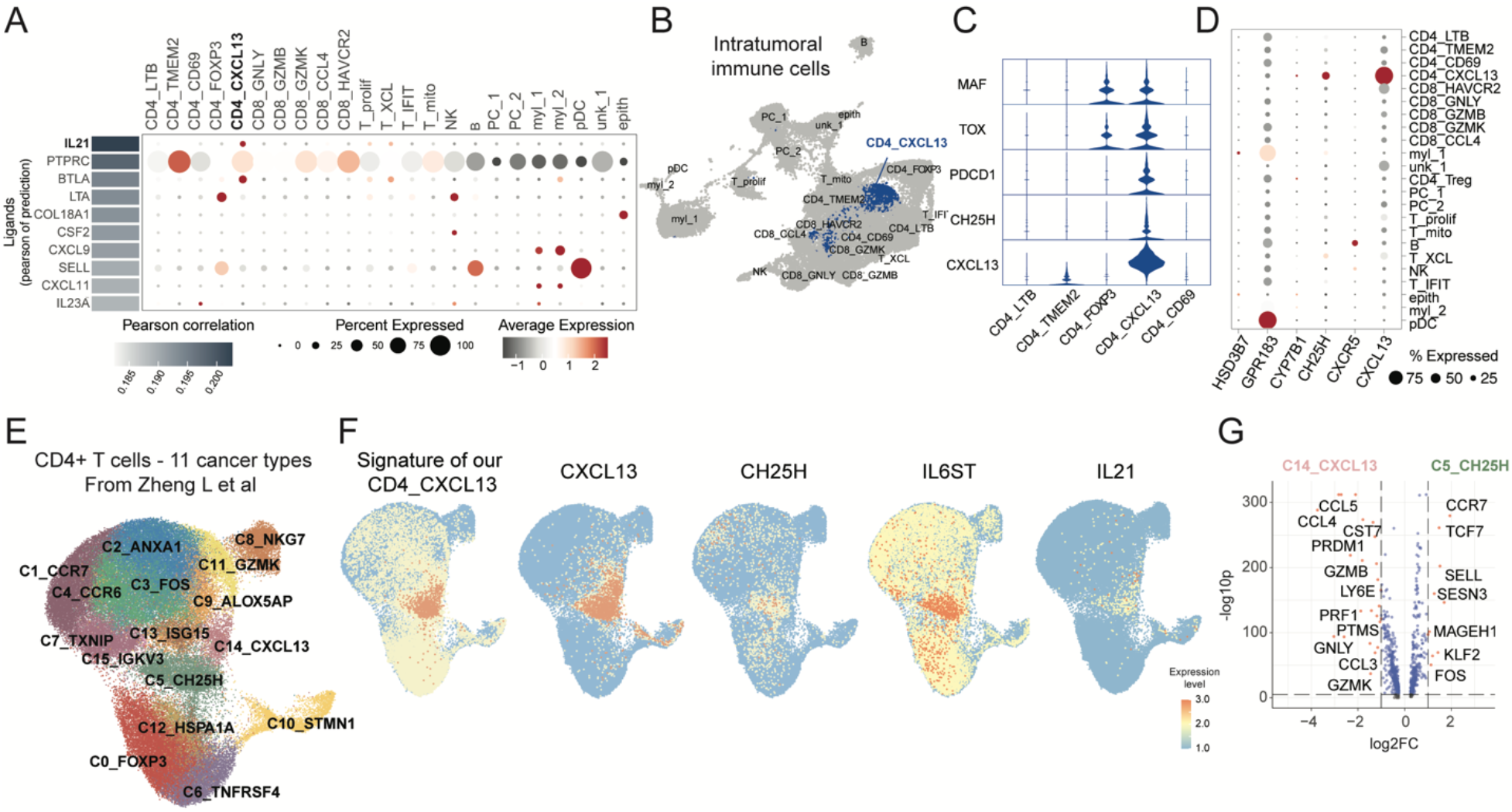
Intratumoral Tfh-like cells overexpress genes involved in immune cell recruitment. Cell communication analysis was performed using the package NicheNet. (**A**) The heatmap shows the top10 ligands promoting interaction with cluster CD8_GZMH, and the dot plot represents their expression on intratumoral immune cells. (**B**) UMAP of intratumoral immune cells of 5 high-grade serous ovarian cancer patients. (**C**) Violin plot showing the expression of some markers of CD4_CXCL13 cluster among CD4 cells. (**D**) Dot plot showing the expression of genes involved in cell recruitment through CXCL13 and 7α,25-HC gradients. (**E**) UMAP of CD4 cells from external dataset (*35*). (**F**) Expression of CD4_CXCL13 signature from our dataset and some of its markers on external CD4 dataset. (**G**) Volcano plot showing the analysis of differentially expressed genes comparing the clusters C5_CH25H and C14_CXCL13.

### Tfh-like cells upregulate genes involved in oxysterol production

Accessing the gene signature of CD4_CXCL13, we identified upregulation of exhaustion-related genes, such as *TOX* and *PDCD1*, the chemokine *CXCL13*, and *CH25H*, a gene encoding cholesterol 25-hydroxylase (Fig. 4C). This enzyme participates in the metabolism of oxysterols by converting cholesterol into 25-hydroxycholesterol (25-HC), which is then processed by CYP7B1 to produce 7α,25-HC, the most active ligand for EBI2 (encoded by the gene *GPR183*) (*43*). In our dataset, plasmacytoid dendritic cells (pDCs) (Fig. 4D) and precursor-exhausted CD8 T cells (Fig. 2C) upregulate *GPR183*, suggesting that besides CXCL13, CD4_CXCL13 cells produces 7α, 25-HC and may recruit other immune subsets to the TME.

To validate the CD4_CXCL13 signature in other cancer types, we analyzed CD4 T cells studied by Zheng L, et al (*35*). This validation cohort represented the CD4 T cell landscape from 11 cancer types and revealed two clusters upregulating *CXCL13*, the C5_CH25H and C14_CXCL13 (Fig. 4E). The cells within C5_CH25H highly scored with cells from our ovarian cancer dataset and upregulated genes identified in CD4_CXCL13 (*CXCL13, CH25H, IL21, TOX, IL6S*, and *PDCD1*) (Fig. 4F, fig. S4A table S4). The frequency of C5_CH25H among CD4 T cells was predominantly higher than C14_CXCL13 (fig. S4B) and usually expressed higher levels of genes *CH25H, CXCL13, IL21*, and *CXCR5* across cancer types (fig. S4C). Differential gene expression analysis between these clusters identified genes related to precursor memory-like phenotype upregulated in C5_CH25H, such as *CCR7, TCF7*, and *SELL*, whereas C14_CXCL13 differentially expressed genes associated with CD8 T cell activation and effector function, including *CCL4, CCL5, CST7, GZMB*, and *PRF1* (Fig. 4G and fig. S4D). Together, our results confirm the presence of cells with CD4_CXCL13 profile in several cancer types and suggest the central role of Tfh-like cells in the formation of lymphoid aggregates in tumors through the secretion of CXCL13 and oxysterol metabolites.

### Tfh-like cells correlate with survival and response to ICB

We next evaluated the prognostic value of aggregates formed by CD4_CXL13 cells in ovarian cancer patients from the TCGA database and in ICB-treated patients using the PredictIO.ca platform (*44*). The genes selected in our signature represent the CD4_CXCL13 cluster (*CD3G, CD4, CXCL13, CH25H*, and *IL21*), the precursor exhausted cells (*CD3G, CD8A, GZMK, TCF7*, and *GPR183*), the plasmacytoid dendritic cells (*CLEC4C* and *GPR183*), and the B cells (*CD79A, MS4A1*, and *CD19*). We tested these signatures either individually (fig. S5) or combined (Fig. 5A) in the TCGA high-grade serous ovarian cancer dataset (352 patients). While Tfh-like and B cell signatures were found to be associated with better survival, no association was observed for precursor-exhausted and pDC signatures. Notably, the combined signatures associated with better prognosis in TCGA and ICB-treated patients, outperforming many of the other published gene expression signatures, including a TLS signature (*18*).

**Fig. 5.**
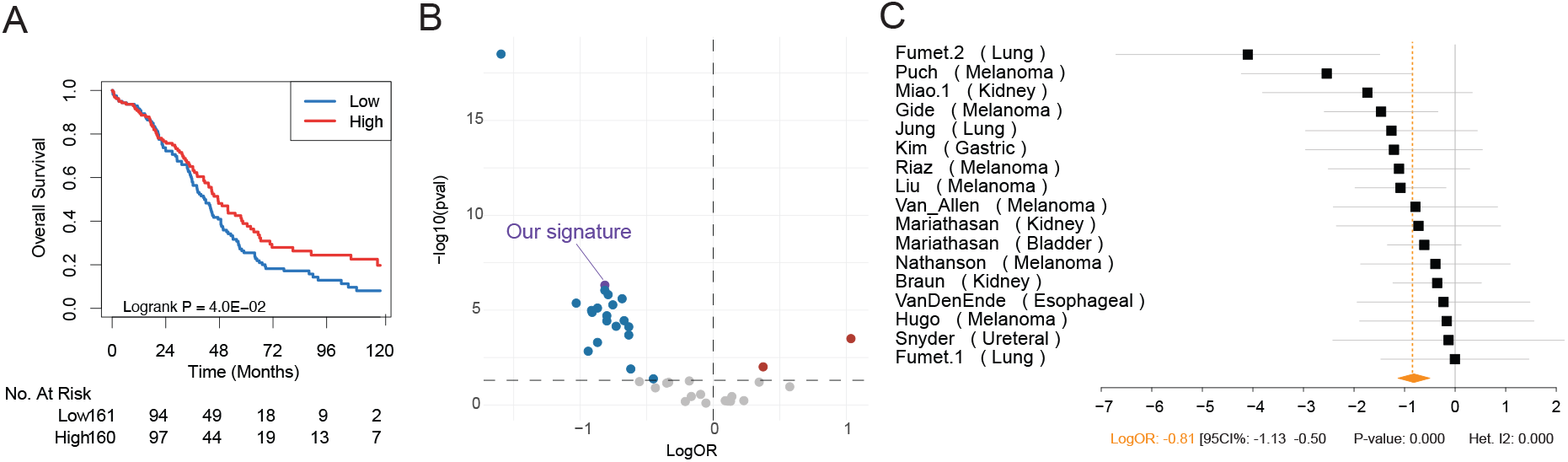
Prognostic impact of signature representing lymphoid aggregates formed by CD4_CXCL13. (A) Kaplan-Meier analysis based on signature representing CD4_CXCL13, precursor-exhausted CD8 T cells, B cells, and pDCs (CD3G, CD4, CXCL13, IL21, CH25H, CD8A, GZMK, TCF7, GPR183, CLEC4C, CD79A, MS4A1, and CD19) in the high-grade serous ovarian cancer cohort from the TCGA study (n=352 patients). (**B**) The impact on response to ICB was evaluated in PredictIO. Meta-analysis of the CD4_CXCL13 gene signature association with ICB response. (**C**) Forest plot displaying for each individual study in black, the log odd-ratios (logOR) and 95% confidence intervals (CI). Orange diamonds represent the logOR and the 95% CI of the meta-analysis.

## DISCUSSION

Although infiltration of immune cells into ovarian tumors is associated with better overall survival (*45*), clinical trials show modest response to ICB (*46*). To understand the antitumor immune response in ovarian cancer, we investigated the dynamics of CD8 T cell subsets in high-grade serous ovarian cancer patients. Studying the transcriptome and TCR repertoire of peripheral and tumor-infiltrating T cells, we identified cell states and inferred their trajectory toward effector function, tissue residency, and exhaustion (schematic representation of the proposed model in fig. S6). Tracking dual-expanded clones and studying cell-cell interactions, we discovered that *IL21* produced by Tfh-like cells (cluster CD4_CXCL13) associated with the expression of genes related to the effector function of CD8 T cells (Fig. 3A and fig. S3B-C). Importantly, Tfh-like cells expressed genes essential for the recruitment of other cells, such as the chemokine *CXCL13* and the enzyme *CH25H*, pointing to their central role in the formation of lymphoid aggregates in the tumor.

Ovarian tumors were enriched with tissue-resident exhausted CD8 T cells. We showed a transitional state of effector exhausted T cells (cluster CD8_GZMH) upregulating *PDCD1, TOX* (Fig. 2B-C), *GZMH*, and *PRF1* (fig. S1D and table S2). This cluster was enriched with dominant clones that were dual-expanded (blood and tumor) (Fig. 3A, C-D) although more frequent in the blood than tumor (Fig. 2A). The trajectory analysis (Fig. 2E) and TCR overlap (Fig. 3B) confirmed the transitional state. We, therefore, propose that cells within the CD8_GZMH cluster reinforce the commitment to tissue residency and exhaustion programs as they progress toward CD8_CCL4, CD8_XCL1, and CD8_CXCL13 states. This transitional state has been recently described in lung tumors (*37*), and we confirmed similarity with our dataset (Fig. 2D). Interestingly, the subsequent states are characterized by clonal expansion and high expression of chemokines, such as CCL4, XCL1, and CXCL13. CCL4 guides the interaction of naïve CD8 T cells with dendritic and CD4 T cells in the lymph node, and disruption in this process compromises the formation of memory CD8 T cell (*47*). CCL4-producing T cells are clonally expanded in tumors (*48*) and peripheral blood of head and neck (HNC) and metastatic melanoma patients (*49*), respectively. XCL1 is a chemoattractant of XCR1+ dendritic cells, important in antigen presentation, activation, and proliferation of CD8 T cells (*50*), and when produced by intratumoral NK cells, DCs are recruited into tumors (*51*). Finally, tumor-specific CD8 T cells are terminally-exhausted and upregulate *ITGAE, ENTPD1*, and the chemoattractant of B cells *CXCL13* (*12, 52*–*54*). Regarding the tissue-residency program in ovarian tumors, the generation and retention of tissue-resident T cells rely on the chemokine receptor CXCR6 (*55*), which was expressed at high levels in the tissue-resident clusters (CD8_ZNF683, CD8_CCL4, and CD8_CXCL13), along with *ITGAE* and *ZNF683*, a gene upregulated in tissue-resident precursor T cells (*56*) also regulating tissue retention of lymphocytes (*57*). In summary, we identified four states that represent clonally-expanded CD8 T cells committed to the exhaustion tissue-resident program, with transcriptional changes to decrease effector function and enhance recruitment and guidance of lymphoid cells in the tumor microenvironment by the production of chemokines (fig. S6).

We furthermore identified two subsets of precursor memory cells defined by the expression of *TCF7, SELL*, and *CCR7* (fig. 2F and G). Because these cells represent a fraction of CD8_ZNF683 and CD8_GZMK, we finely discriminated CD8 T cell clusters into subclusters, which k_8 and k_10 within CD8_ZNF683 and k_22, k_16, k_5, and k_3 within CD8_GZMK (fig. S2). While the CD8_GZMK lineage defined cells expressing the memory-like markers described earlier and gradual increase on *GZMH* and *CCL4* levels, the CD8_ZNF683 is separated into two very distinct subclusters, k_8 defined by memory-like markers, and k_10 upregulating the tissue retention markers CXCR6 and ITGAE (Fig. 2F-G). Although very different, those subclusters have large TCR overlaps, corroborating their proximal position in the trajectory analysis (fig. S2D). In fact, most of the shared TCRs of k_8 are with tissue-resident subclusters (k_10, k_11, and k_7) of CD8_ZNF683, CD8_XCL1, and CD8_CXCL13. On the other hand, the few cells expressing the highest levels of TCF7, SELL, and CCR7, are in k_22 (within CD8_GZMK), which also upregulate the inhibitory receptors *TIGIT, HACR2*, and *CTLA4* (fig. S2B-C). Interestingly, this profile is similar to one of the two subsets of memory-like precursor T cells identified in humans (*58*). While one of those precursors differentiates into effector T cells, the other was characterized as precursor-exhausted CD8 T cells upregulating *GZMK, PDCD1*, and *TIGIT*, such as the k_22 in our study. Therefore, the memory-like precursor cells identified in our study are divided into those similar to tissue-resident T cells and those within the GZMK lineage. In both cases, the cells upregulate genes essential for recirculation, including SELL, CCR7, and KLF2 (Fig. 2F-G). Thus, we proposed the existence of two subsets of recirculating precursor memory cells, where one relates to tissue-resident cells and the other to the GZMK lineage.

By including peripheral blood and tumor samples, we could identify states of recirculating memory cells and dual-expanded effector cells. Considering the importance of dual-expanded clones in the antitumor immune response (*14*), we interrogated the cellular interactions they encountered in the tumor microenvironment. Using the tool NicheNet, we identified *IL21* as one of the top ligands mediating the interaction between dual-expanded clones and CD4_CXCL13 (Tfh-like cells) (Fig. 4A). Interestingly, the gene signature of Tfh-like cells showed that besides dual-expanded clones, these cells might attract other immune cells through CXCL13 and oxysterol metabolites. We identified B cells and pDCs as the major cell types recruited by the CXCL13-CXCR5 axis and oxysterol-GPR183, respectively. Additionally, the CD8_GZMK also upregulate *GPR183*, suggesting that a gradient of oxysterol might recruit them to a region defined by Tfh-like cells (Fig. 2C). Importantly, high expression of *GZMK* is observed in a distinct subset of CD8 T cells, showed expanded in aging (*59*), identified as precursor-exhausted T cell in healthy donor blood (*58*), associated with excluded ovarian tumor (T cell restricted to tumor stroma) (*60*), and identified in breast cancer as pre-effector/memory cells associated with low T expansion following anti-PD-1 treatment (*40*). Together with our results, these studies suggest that GZMK_CD8 T cells require additional signals to differentiate into effector CD8 T cells, which may occur in a specific niche in the tumor microenvironment defined by CD4_CXCL13. Tfh-like cells in non-lymphoid tissues support humoral and cell-mediated responses by recruiting B cells and supplying IL21 (*21, 23, 26, 27, 30, 31, 42, 61*). Indeed, a recent study showed that Tfh-like cells recruit precursor CD8 T cells and DCs using the oxysterol metabolite 7α,25-HC, which may support a niche for differentiation of effector cells in hepatocellular carcinomas (*62*). Moreover, our meta-analysis on the TCGA cohort of HGSOC and several cohorts of ICB-treated showed the improved clinical response associated with these lymphoid aggregates formed by Tfh-like cells.

We here used gene expression, TCR repertoire, and cell interaction analysis to understand immunity cycles in ovarian cancer. Studying peripheral and intratumoral lymphocytes, we identified cellular programs driven in the tumor microenvironment (exhaustion, tissue retention, and effector function) that may require a specific niche formed by Tfh-like cells (schematic representation in fig. S6). We demonstrated two subsets of recircuiting precursor CD8 T cells, one related to tissue-resident clones (CD8_ZNF683) and the other identified as pre-effector cells (CD8_GZMK). The former may require the niche formed by Tfh-like cells to proliferate and differentiate into effector cells, which may help understand the association of CD8_GZMK cells with excluded tumors, poor T cell expansion, and the prognostic value of TLSs. We proposed that Tfh-like cells use oxysterol to recruit CD8_GZMK and plasmacytoid dendritic cells into tumors, a mechanism well described in SLO but poorly explored in tumor immunity. Understanding the formation of lymphoid aggregates in humans remains a challenge. Collectively, our results suggest that Tfh-like cells expressing IL-21 help promote antitumor immunity against ovarian tumors by coordinating the action of immune cells responsive to 7α-25-HC.

## MATERIALS AND METHODS

### Human specimens

This study was approved by the Research and Medical ethical committee of Montreal University Hospital Center (CÉR-CHUM). All participants signed the written informed consent and had their tissue banked at CRCHUM Ovarian Cancer biobank. Fresh tissue was collected from treatmentnaive patients undergoing debulking surgery for ovarian cancer and immediately processed to obtain a single-cell suspension. Pathologists confirmed the ovarian cancer subtype, and cases classified as high-grade serous carcinoma were included in this study. Following the manufacturer’s instructions, the tumor was dissociated using the mechanical and enzymatic system GentleMACS(tm) (MACS Miltenyi). Viable cells (Dead Cell Removal kit, MACS Miltenyi) and immune cells (human TIL-CD45 microbeads, MACS Miltenyi) were isolated and cryopreserved in 90% FBS and 10% DMSO.

### Preparation of scRNA-seq libraries

CD45+ cells were characterized at the single-cell level using Chromium technology (10X Genomics). Version 1.1 of Next GEM Single Cell 5’ Library kit revealed gene expression and TCR repertoire of CD45+ cells isolated from the tumor (5 patients) and matched blood (4 patients and two healthy donors). Cells were thawed in FBS, treated with 200U/mL of Nuclease S7, washed three times, and resuspended in 0.04% BSA PBS 1X to a final concentration of 800-1000 cells/μL. Cell viability was assessed by Trypan Blue stain, and samples with at least 80% viability were loaded into a 10X system for RNA barcoding. The libraries were sequenced on Illumina NovaSeq 6000 targeting a minimum sequencing depth of 50,000 read pairs per cell (26 bp read 1, 8bp i7 index, 91bp read 2) in the gene expression library and a minimum of 5,000 reads per cell (26 bp read 1, 8bp i7 index, 91bp read 2) in TCR libraries. The sequencing was performed at the Centre for Pediatric Clinical Genomics of CHU Sainte-Justine, Montreal, CA.

### Processing of scRNA-seq data

The scRNA-seq libraries were processed using the software Cell Ranger (10X Genomics) version 3.1. Illumina’s output files Base Call (BCL) were demultiplexed, and fastq files were generated using the command’ cellranger mkfastq’. The gene expression matrix was generated with the command’ cellranger count’ using the human genome reference GRCh38. The filtered matrix was used for further analysis. We recovered 43,948 and 40,303 cells from the tumor and blood tissue. From 24,061 recovered genes, an average of 1,264 genes were detected per cell. The TCR library was processed using the ‘cellranger vdj’ command. The filtered contig annotation was used for T cell clonotype analysis. Detailed results on the quality of sequencing are available in table S1.

### Analysis of scRNA-seq data on intratumoral CD45+ cells

Analysis was performed on Seurat package v3.2 (*63*). Cells were filtered by the number of genes detected (>200 and <6000 genes) and expressed genes were defined by expressed in at least 3 cells. Subsequently, dead cells (>15% mapped to the mitochondrial genome) or cancer cells (expression of MUC16 and EPCAM) were filtered out. The dataset was normalized by the scTranform method (*64*), followed by regressing out the mitochondrial and heat shock (obtained in HUGO Gene Nomenclature Committee) genes expression. Before the integration of the five patients’ data, 3000 features were selected using ‘SelectIntegrationFeatures’ to run ‘PrepSCTIntegration’ further. These features were used to identify anchors and integrate the data using the functions ‘FindIntegrationAnchors’ and ‘IntegrateData’, respectively. For data visualization, dimensionally reduction was executed using the functions RunPCA and RunUMAP (in the first 60 principal components), following clustering by FindNeighbors (in the first 60 principal components) and ‘FindClusters’. Clusters were characterized by differentially expressed genes analysis using the test Model-based Analysis of Single-cell Transcriptomics (MAST) (*65*) in the function ‘FindAllMarkers’. Cell type annotation was performed manually and then confirmed by the automatic method SingleR (*66*). The top genes identified by the previous analysis, as well as known markers, were used to identify cell types: cytotoxic T cells (*CD3D, CD3G, CD8A*), helper T cells (*CD3D, CD3G, CD4*), T regulatory cells (*CD4, FOXP3*), proliferating lymphocytes (*MKI67*), B cells (*MS4A1, CD19, CD79A*), Natural Killer cells (*NCAM1, KLRC1*), macrophages (*CD68, FCGR3A, CD14*), dendritic cells (*CLEC4C, CST3*), endothelial cells (*VWF*), epithelial cells (*EPCAM*).

### Analysis of scRNA-seq on peripheral CD45+ cells

The blood dataset was analyzed as the tumor dataset, except that the filtering of cancer cells and regression of heat shock genes in normalization were not performed.

### Analysis of T cell dataset

To reveal the heterogeneity of T cells in tumor and blood of HGSOC patients, clusters of T cells identified in cancer and blood datasets were selected for further integration following scTransform in the Seurat pipeline. We calculated doublets with the tools’ DoubletFinder’ (*67*), ‘Scrublet’ (*68*), and ‘DoubletDecon’ (*69*). Regarding TCR-seq, the clones of T cells were classified as CellRanger software. We obtained this information in the output file “filtered_contig_annotations.csv,” Since the clonotype is classified within a sample, we had to rename them to guarantee that clones would not be named as in other samples. We used the TCRαβ repertoire to determine the diversity, clonal expansion, and clonal overlap. For diversity, we used the diversity 50 index (D50), which corresponds to the percentage of dominant T cell clones accounting for the cumulative 50% of total CDR3 sequences. The clonal overlap was calculated using the Morisita-Horn index (MHI) in the R package ‘divo’ (*70*). For clonal expansion, the clones were classified into five groups according to their expansion pattern as proposed in Wu et all (*14*): represented by one cell within a tissue (singleton), multiple cells within a tissue (multiplets), or one or more cells in both tissue (dual-expanded). We used the databases TBAdb (*71*), McPAS-TCR (*72*), and VDJdb (*73*) to identify T cells specific to known antigens.

Four clusters represented either cell types we couldn’t assign or non-T cells (‘myl’, ‘uk_1’, ‘uk_2’, ‘uk_3’). To characterize those clusters, we looked at the frequency of TCR sequenced, the frequency of doublets, and the level of expression of *CD3G*. The ratios of doublets were evenly distributed in all clusters, suggesting that those clusters do not represent more than one cell captured in the scRNA-seq assay. The cluster myl is probably myeloid cells contaminating our dataset because of reduced αβ TCR-seq, low expression of CD3G (Supplementary Fig. 1B), and upregulation of the genes involved in the antimicrobial response of myeloid cells (*S100A8, S100A9*, and *LYZ*) (Suppl fig 1F). The cluster uk1 upregulates *TRBV11-2*, which has been recently reported expanded in patients with severe multisystem inflammatory syndrome in children (MIS-C) (*74*). The cluster uk2 groups cells upregulating long non-coding RNAs (*RP11-290D2*.*6* and *RP11-124N14*.*3*), while uk3 upregulates genes involved in antigen presentation by MHC class II.

### Cell communication and trajectory analysis

The package NicheNet 1.1.0 version (*41*) was used to reveal the interactions of dual-expanded clones within the tumor immune microenvironment (TIME). This approach predicted the ligand-receptor network based on genes differentially expressed in intratumoral compared to peripheral dual-expanded clones in clusters CD8_GZMH and CD8_GZMB. Trajectory analysis with PAGA was performed following Scanpy version 1.6.0 (*75*) pipeline in Python version 3.8.3. We exported the Seurat integrated assay of 2000 variable genes to the h5 Anndata file, denoised the graph computing diffusion map, and computed PAGA on clusters identified by the Leiden algorithm.

### External data

CD8 T cells signatures from other studies were accessed in the supplementary data of Gueguen et al (table S2 (*37*)), Guo et al (table S3 (*34*)), Wu et al (table S4 and S5 (*14*)). The CD4 dataset of Zheng L et al (*35*) was obtained in the zenodo repository (DOI 10.5281/zenodo.5461803) and analyzed in Seurat using Harmony integration (*76*).

### Association with patient outcome

ICB studies were curated as described in Bareche et al. (*44*). Meta-analysis was performed on the PredictIO.ca web-application. Briefly, the Association of specific biomarkers with ICB response was assessed using a logistic regression model. To improve reproducibility, the results of individual independent studies were pooled using random-effects meta-analysis with inverse variance weighting in DerSimonian and Laird random-effects models (*77*). Heterogeneity across studies was evaluated by using the Q statistic along with I^2^ index, which describes the total variation across studies attributable to heterogeneity rather than sampling error (*78*–*80*). Note that I^2^ value of greater than 50% along with Cochran’s Q statistic P < 0.05 represent moderate to high heterogeneity (*78*). Subgroup analysis was considered to assess the impact of tumor type, sequencing technology and normalization method on the source of moderate to high heterogeneity (*77*). Potential publication bias was performed using the funnel plot and the Egger test. No statistically significant publication bias was observed (data not shown). All analyses were performed on the R platform (version 3.6.3). When needed, p-values were corrected for multiple testing using the Benjamini-Hochberg (False Discovery Rate, FDR) method. Associations were deemed statistically significant for p-values and/or FDR lower or equal to 0.05.

## SUPPLEMENTARY MATERIALS

Fig. S1. Strategy to define T cell subsets in high-grade serous ovarian cancer.

Fig. S2. Characterization of subclusters of CD8 T cells.

Fig. S3. Genes modulated by cellular communication between dual-expanded effector CD8 T cells and the tumor immune microenvironment (TIME).

Fig. S4. Markers of CD4_CXCL13 conserved in several cancer types.

Fig. S5. Impact of Tfh-like, B, pDC, and CD8 T memory-like cell signatures on ovarian cancer survival.

Fig. S6. Model proposed to explain the lineage relationship between intratumoral and peripheral CD8 T cell subsets in HGSOC.

Table S1. Sequencing parameters. Table S2. Markers of T cell clusters.

Table S3. Markers of subclusters of T cells. Table S4. Markers of intratumoral immune cells.

Table S5. Markers of CD4_CXCL13 conserved in several cancer types. Table S6. External signatures.

## FUNDING

Terry Fox Translational Program Grant to the Immunotherapy Network (iTNT) (RL, JS, AMMM, JFC).

Canada Research Chair in Functional Genomics of Reproduction and Development (950-233204) (SP)

Swedish Research Council (2016–01919) (SP, CZ),

Swedish Society for Medical Research (Dnr4-236-2107), (SP, CZ).

Fonds de Recherche du Québec – Santé (MSC, YB)

## AUTHOR CONTRIBUTIONS

Conceptualization: MSC, YB

Methodology: MSC, YB, CZ

Investigation: MSC, YB, CZ, PT, KR

Visualization: MSC, YB, CZ

Resources: AMMM, DP, VS, BC

Funding acquisition: RL, JS, AMMM, JFC

Supervision: RL, JS, SP

Writing – original draft: MSC

Writing – review & editing: MSC, YB, CZ, SP, RL, JS

## COMPETING INTERESTS

JS is a SAB member and owns stocks of Surface Oncology. All other authors declare that they have no competing interests.

## DATA AND MATERIALS AVAILABILITY

The single cell RNA sequencing will be deposited in the Gene Expression Omnibus database and the codes used in the analysis in Bitbucket repository.

