## Supplemental figures for "Predicting developmental relationships of tumor resident and circulating T cells in ovarian cancer"

### Supplementary figures

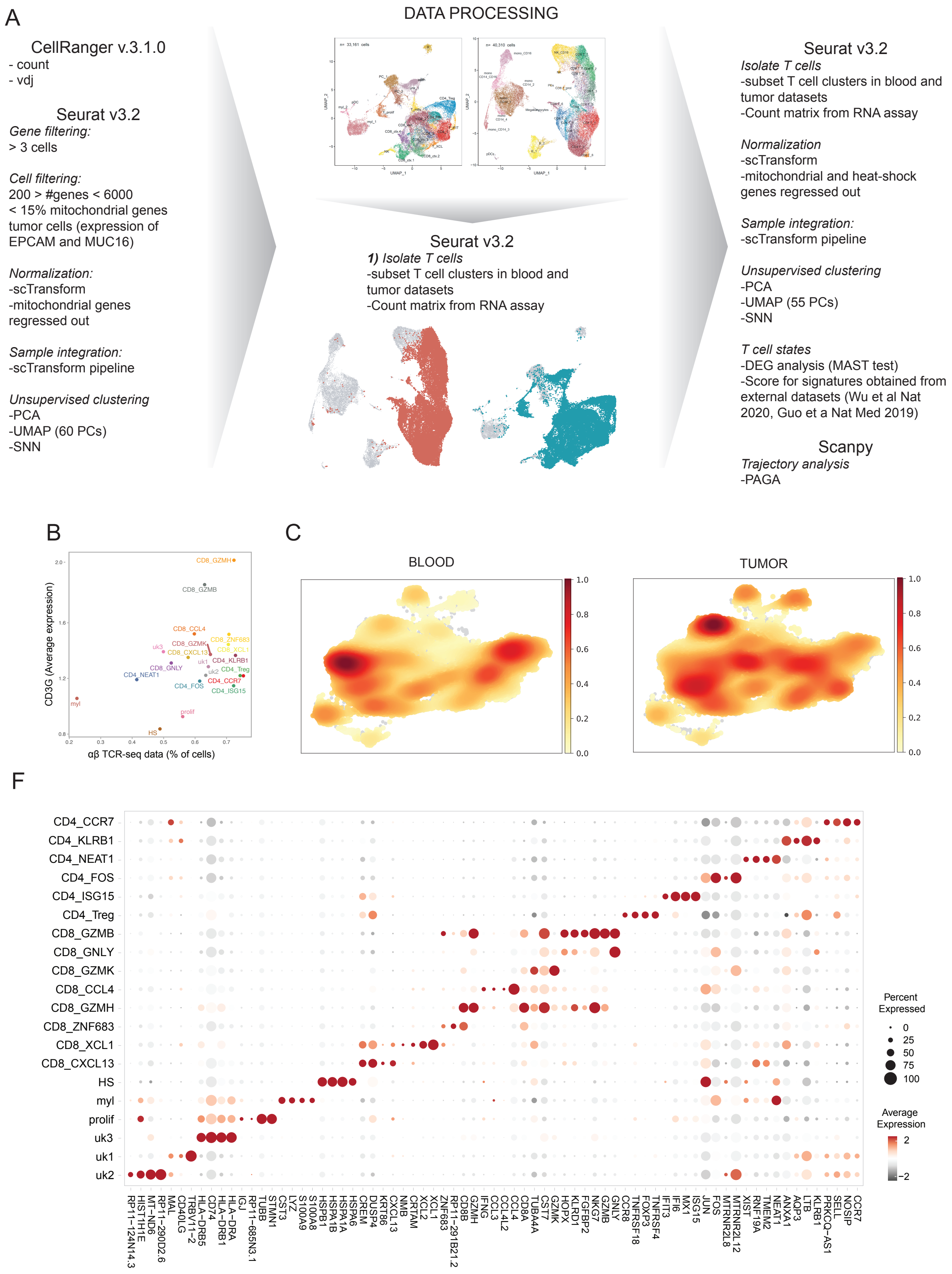
A
DATA PROCESSING
CellRanger v.3.1.0
- count
- vdj
Seurat v3.2
Gene filtering:
> 3 cells
Cell filtering:
200 > #genes < 6000
< 15% mitochondrial genes
tumor cells (expression of EPCAM and MUC16)
Normalization:
-scTransform
-mitochondrial genes regressed out
Sample integration:
-scTransform pipeline
Unsupervised clustering
-PCA
-UMAP (60 PCs)
-SNN
Seurat v3.2
1) Isolate T cells
-subset T cell clusters in blood and tumor datasets
-Count matrix from RNA assay
Seurat v3.2
Isolate T cells
-subset T cell clusters in blood and tumor datasets
-Count matrix from RNA assay
Normalization
-scTransform
-mitochondrial and heat-shock genes regressed out
Sample integration:
-scTransform pipeline
Unsupervised clustering
-PCA
-UMAP (55 PCs)
-SNN
T cell states
-DEG analysis (MAST test)
-Score for signatures obtained from external datasets (Wu et al Nat 2020, Guo et a Nat Med 2019)
Scanpy
Trajectory analysis
-PAGA
B
CD3G (Average expression)
αβ TCR-seq data (% of cells)
CD8_GZMH
CD8_GZMB
CD8_CCL4
CD8_ZNF683
CD8_GZMK
CD8_XCL1
CD8_CXCL13
CD4_KLRB1
CD8_GNLY
CD4_Treg
CD4_NEAT1
CD4_CCR7
CD4_FOS
CD4_ISG15
uk1
uk2
uk3
myl
prolif
HS
C
BLOOD
TUMOR
1.0
0.8
0.6
0.4
0.2
0.0
F
CD4_CCR7
CD4_KLRB1
CD4_NEAT1
CD4_FOS
CD4_ISG15
CD4_Treg
CD8_GZMB
CD8_GNLY
CD8_GZMK
CD8_CCL4
CD8_GZMH
CD8_ZNF683
CD8_XCL1
CD8_CXCL13
HS
myl
prolif
uk3
uk1
uk2
Percent Expressed
0
25
50
75
100
Average Expression
2
−2

**Fig. S1. Strategy to define T cell subsets in high-grade serous ovarian cancer.** (**A**) Schematic representation of scRNA-seq analysis of T cell dataset from 5 high-grade serous ovarian carcinoma patients and two healthy donors. (**B**) Scatter plot of CD3G expression level and proportion of cells for which an αβTCR sequence was detected in each cluster. Bar plots indicate the relative proportion of cells from each patient within a cluster (**C**) and the proportion of cells from each cluster within a patient (**D**). (**E**) UMAP representation of cell abundance in each cluster in blood and tumor. (**F**) Dot plot of top 4 genes identified by differentially expressed gene analysis in each cluster.

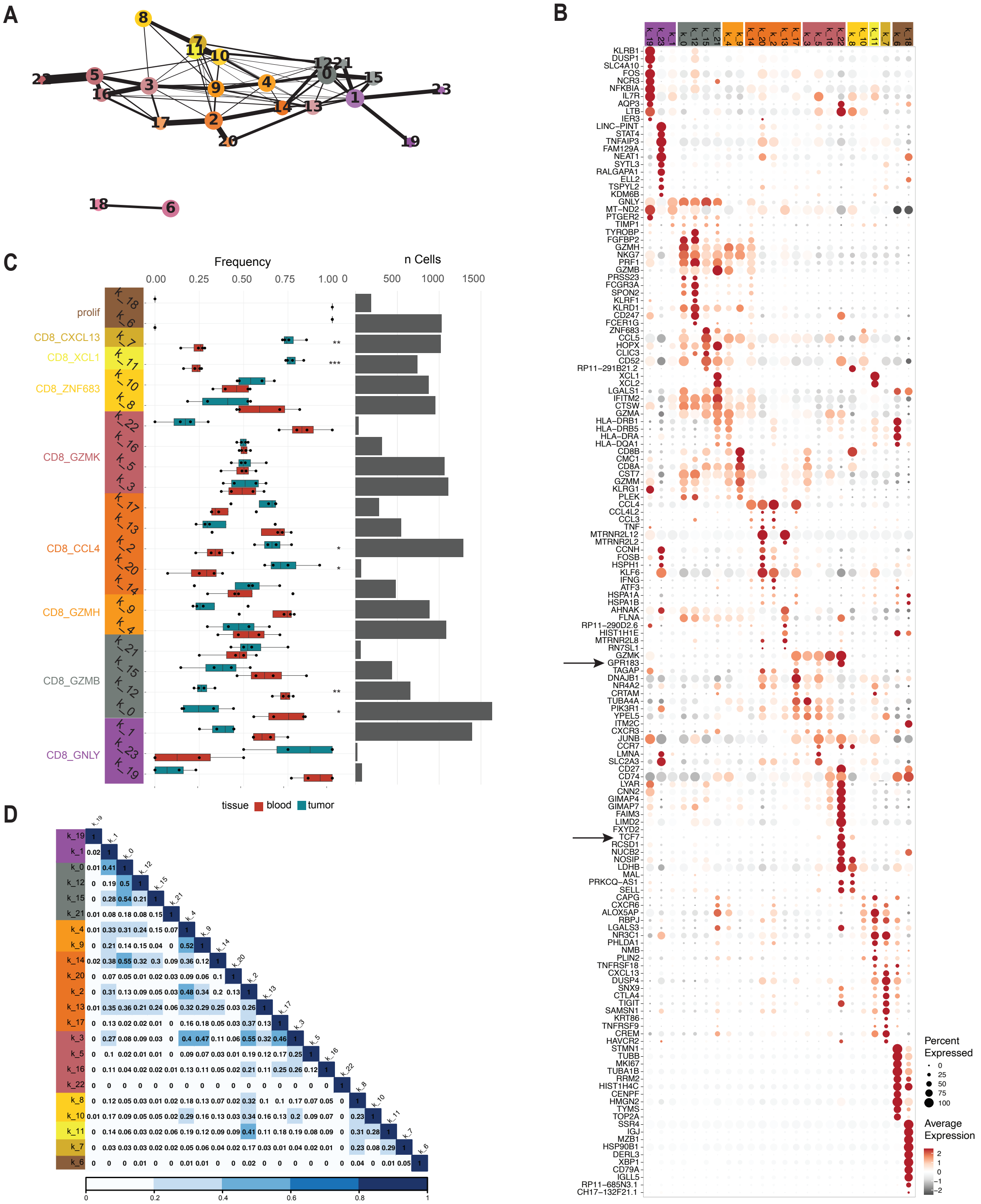
A
B
C
D
Frequency
0.00 0.25 0.50 0.75 1.00
n Cells
0 500 1000 1500
prolif
CD8_CXCL13
CD8_XCL1
CD8_ZNF683
CD8_GZMK
CD8_CCL4
CD8_GZMH
CD8_GZMB
CD8_GNLY
tissue blood tumor
Percent Expressed
0 25 50 75 100
Average Expression
2 1 0 −1 −2
KLRB1
DUSP1
SLC4A10
FOS
NCR3
NFKBIA
IL7R
AQP3
LTB
IER3
LINC−PINT
STAT4
TNFAIP3
FAM129A
NEAT1
SYTL3
RALGAPA1
ELL2
TSPYL2
KDM6B
GNLY
MT−ND2
PTGER2
TIMP1
TYROBP
FGFBP2
GZMH
NKG7
PRF1
GZMB
PRSS23
FCGR3A
SPON2
KLRF1
KLRD1
CD247
FCER1G
ZNF683
CCL5
HOPX
CLIC3
CD52
RP11−291B21.2
XCL1
XCL2
LGALS1
IFITM2
CTSW
GZMA
HLA−DRB1
HLA−DRB5
HLA−DRA
HLA−DQA1
CD8B
CMC1
CD8A
CST7
GZMM
KLRG1
PLEK
CCL4
CCL4L2
CCL3
TNF
MTRNR2L12
MTRNR2L2
CCNH
FOSB
HSPH1
KLF6
IFNG
ATF3
HSPA1A
HSPA1B
AHNAK
FLNA
RP11−290D2.6
HIST1H1E
MTRNR2L8
RN7SL1
GZMK
GPR183
TAGAP
DNAJB1
NR4A2
CRTAM
TUBA4A
PIK3R1
YPEL5
ITM2C
CXCR3
JUNB
CCR7
LMNA
SLC2A3
CD27
CD74
LYAR
CNN2
GIMAP4
GIMAP7
FAIM3
LIMD2
FXYD2
TCF7
RCSD1
NUCB2
NOSIP
LDHB
MAL
PRKCQ−AS1
SELL
CAPG
CXCR6
ALOX5AP
RBPJ
LGALS3
NR3C1
PHLDA1
NMB
PLIN2
TNFRSF18
CXCL13
DUSP4
SNX9
CTLA4
TIGIT
SAMSN1
KRT86
TNFRSF9
CREM
HAVCR2
STMN1
TUBB
MKI67
TUBA1B
RRM2
HIST1H4C
CENPF
HMGN2
TYMS
TOP2A
SSR4
IGJ
MZB1
HSP90B1
DERL3
XBP1
CD79A
IGLL5
RP11−685N3.1
CH17−132F21.1

**Fig. S2. Characterization of subclusters of CD8 T cells.** (**A**) PAGA projection of T cells dataset showing the subclusters. The color is a reference to the cluster most represented within a subcluster. (**B**) Dot plot of top 10 markers for each cluster identified by DEG analysis. (**C**) Bar plots indicate on the left the frequency of cells from blood or tumor in each subcluster and on the right the number of cells in each cluster. Data are expressed as mean ± SD of an individual patient. Significant p values are determined by the t statistics (* $p<0.05$, ** $p<0.01$, *** $p<0.001$). (**D**) Heat map of TCR overlaps calculated by the Morisita-Horn index.

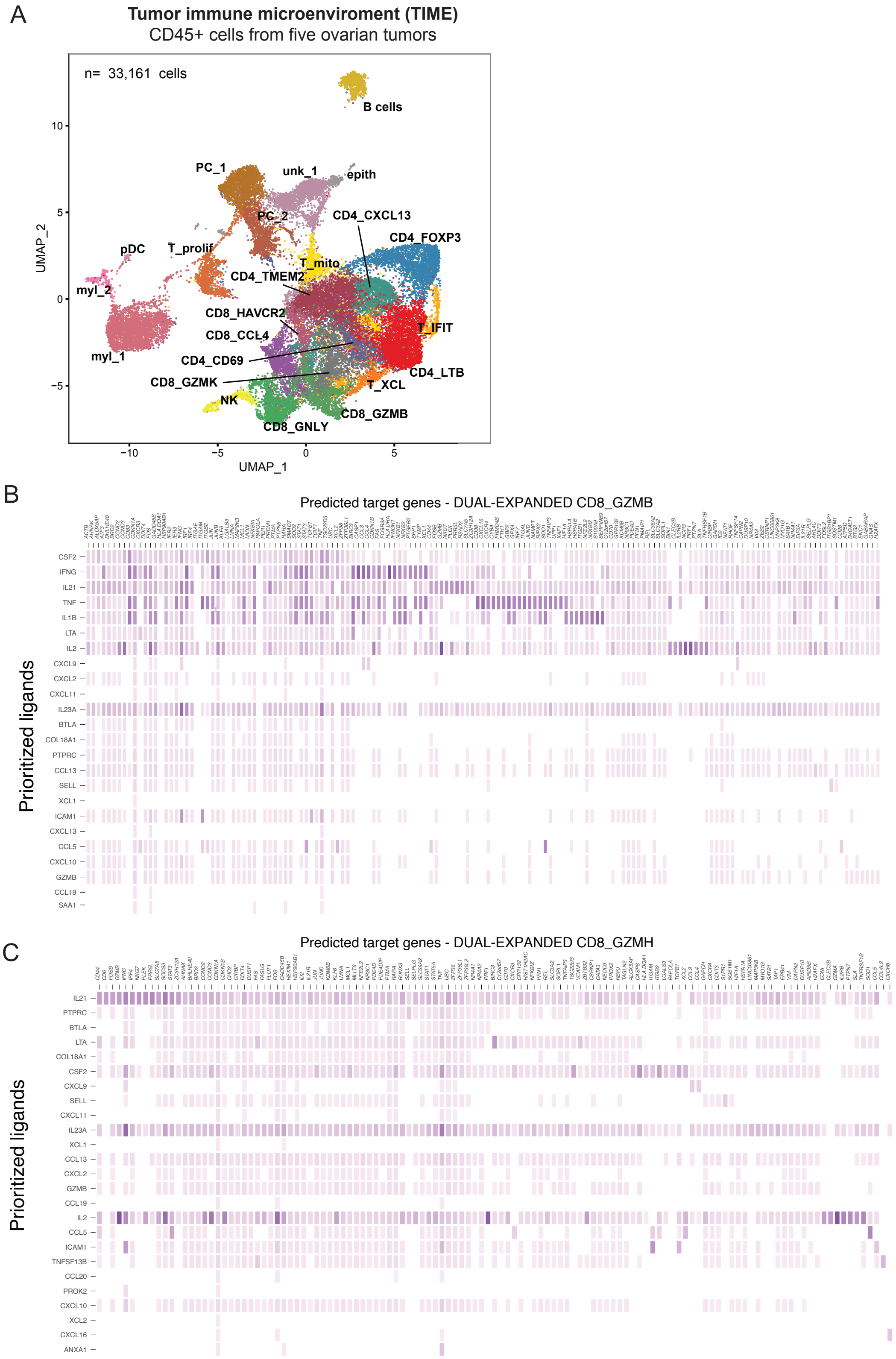
A
Tumor immune microenviroment (TIME)
CD45+ cells from five ovarian tumors
n= 33,161 cells
B cells
PC_1
unk_1
epith
PC_2
CD4_CXCL13
CD4_FOXP3
pDC
T_prolif
T_mito
CD4_TMEM2
myl_2
CD8_HAVCR2
T_IFIT
CD8_CCL4
myl_1
CD4_CD69
CD4_LTB
CD8_GZMK
T_XCL
NK
CD8_GZMB
CD8_GNLY
UMAP_1
UMAP_2
10
5
0
−5
−10
B
Predicted target genes - DUAL-EXPANDED CD8_GZMB
Prioritized ligands
CSF2
IFNG
IL21
TNF
IL1B
LTA
IL2
CXCL9
CXCL2
CXCL11
IL23A
BTLA
COL18A1
PTPRC
CCL13
SELL
XCL1
ICAM1
CXCL13
CCL5
CXCL10
GZMB
CCL19
SAA1
C
Predicted target genes - DUAL-EXPANDED CD8_GZMH
Prioritized ligands
IL21
PTPRC
BTLA
LTA
COL18A1
CSF2
CXCL9
SELL
CXCL11
IL23A
XCL1
CCL13
CXCL2
GZMB
CCL19
IL2
CCL5
ICAM1
TNFSF13B
CCL20
PROK2
CXCL10
XCL2
CXCL16
ANXA1

**Fig. S3. Genes modulated by cellular communication between dual-expanded effector CD8 T cells and the tumor immune microenvironment (TIME).** We inferred cell-cell interactions between dual-expanded effector CD8 T cells (within CD8_GZMB and CD8_GZMH clusters) and other immune cells within ovarian tumors. (**A**) The UMAP projection represents the immune cells (CD45+ cells) composing tumors of 5 ovarian cancer patients. (**B and C**) The heatmap represents the top25 ligands expressed by sender cells (the cells composing the TIME) and their regulatory potential in target genes identified differentially expressed in intratumoral versus peripheral dual-expanded in the clusters CD8_GZMH (**B**) and CD8_GZMB (**C**).

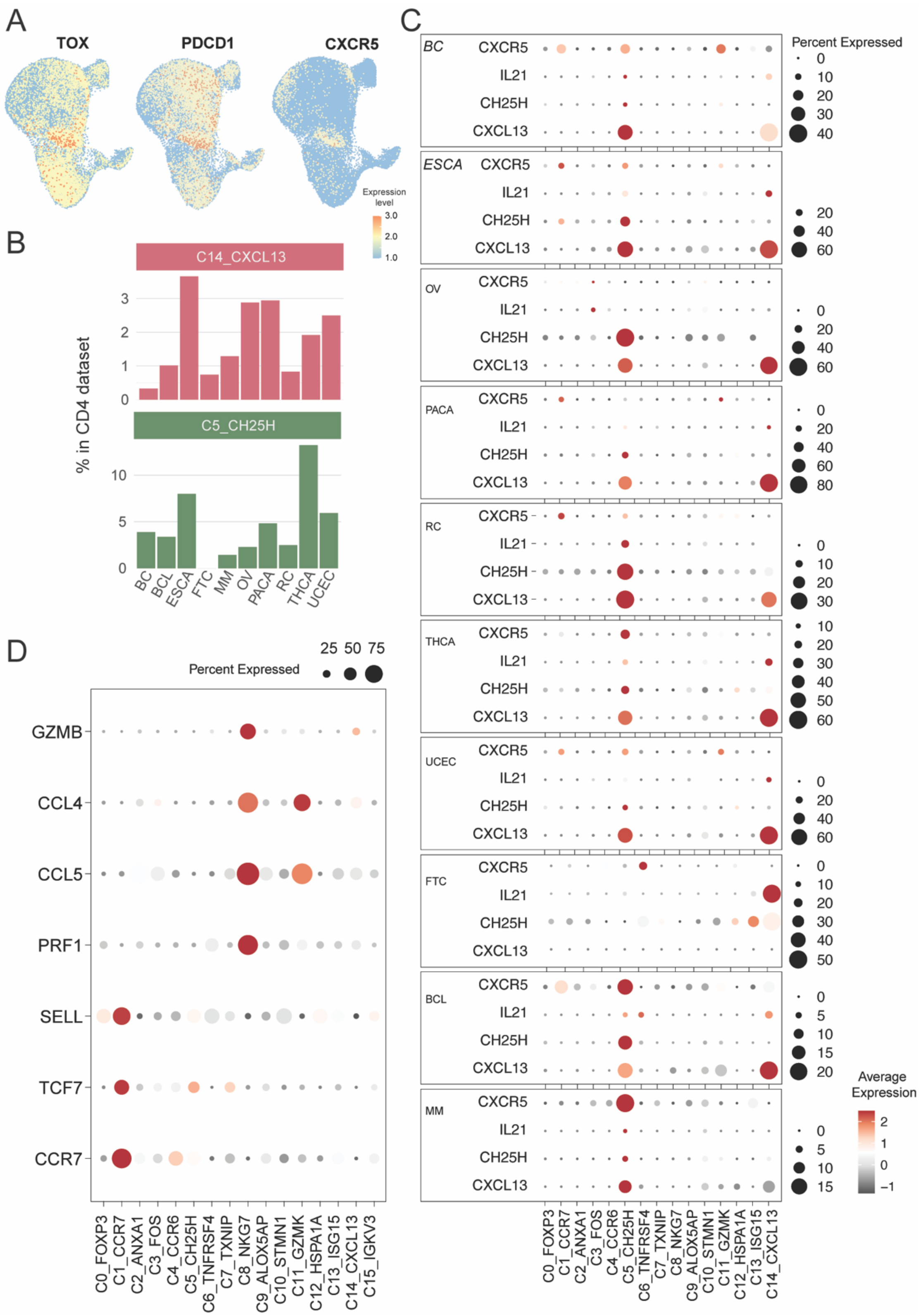
A
TOX
PDCD1
CXCR5
Expression level
3.0
2.0
1.0
B
C14_CXCL13
C5_CH25H
% in CD4 dataset
BC
BCL
ESCA
FTC
MM
OV
PACA
RC
THCA
UCEC
C
BC
ESCA
OV
PACA
RC
THCA
UCEC
FTC
BCL
MM
CXCR5
IL21
CH25H
CXCL13
Percent Expressed
Average Expression
C0_FOXP3
C1_CCR7
C2_ANXA1
C3_FOS
C4_CCR6
C5_CH25H
C6_TNFRSF4
C7_TXNIP
C8_NKG7
C9_ALOX5AP
C10_STMN1
C11_GZMK
C12_HSPA1A
C13_ISG15
C14_CXCL13
D
Percent Expressed
25 50 75
GZMB
CCL4
CCL5
PRF1
SELL
TCF7
CCR7
C15_IGKV3

**Fig. S4. Markers of CD4_CXCL13 conserved in several cancer types.** (**A**) UMAP represents the expression level of some exhaustion-related genes (*TOX* and *PDCD1*) and the Tfh markers *CXCR5* in the external/validation dataset. (**B**) Bar plot indicating the frequency of clusters C5_CH25H and C14_CXCL13 within CD4 cells across several cancer types. (**C**) Dot plots showing the expression level of genes upregulated in our ovarian cancer dataset in the external/validation dataset across cancer types. (**D**) Dot plot showing the expression level of top genes identified in the DEG of C5_CH25H compared to the o C14_CXCL13 cluster. BC: Breast cancer, ESCA: esophageal cancer, OV: ovarian cancer, PACA: pancreatic cancer, RC: renal carcinoma, THCA: thyroid carcinoma, UCEC: uterine corpus endometrial carcinoma, FTC: fallopian tube carcinoma, BCL: B-cell lymphoma, MM: multiple myeloma.

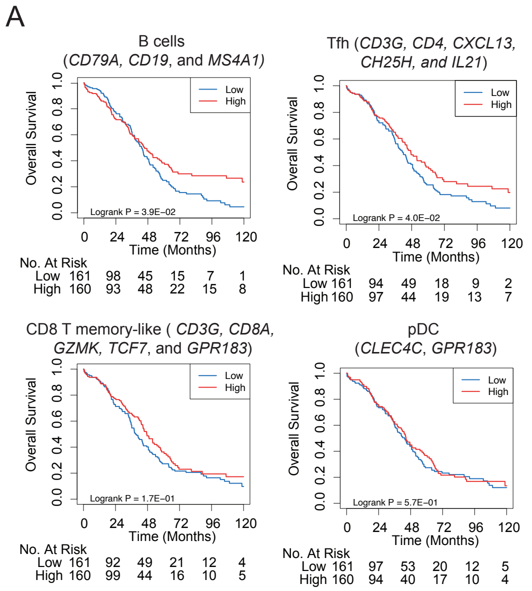


**Fig. S5. Impact of Tfh-like, B, pDC, and CD8 T memory-like cell signatures on ovarian cancer survival.** (**A**) Kaplan-Meier analysis in the high-grade serous ovarian cancer cohort of TCGA study based on signatures of B cells, Tfh-like cells, precursor-exhausted CD8 T cells, and plasmacytoid dendritic cells (pDCs).

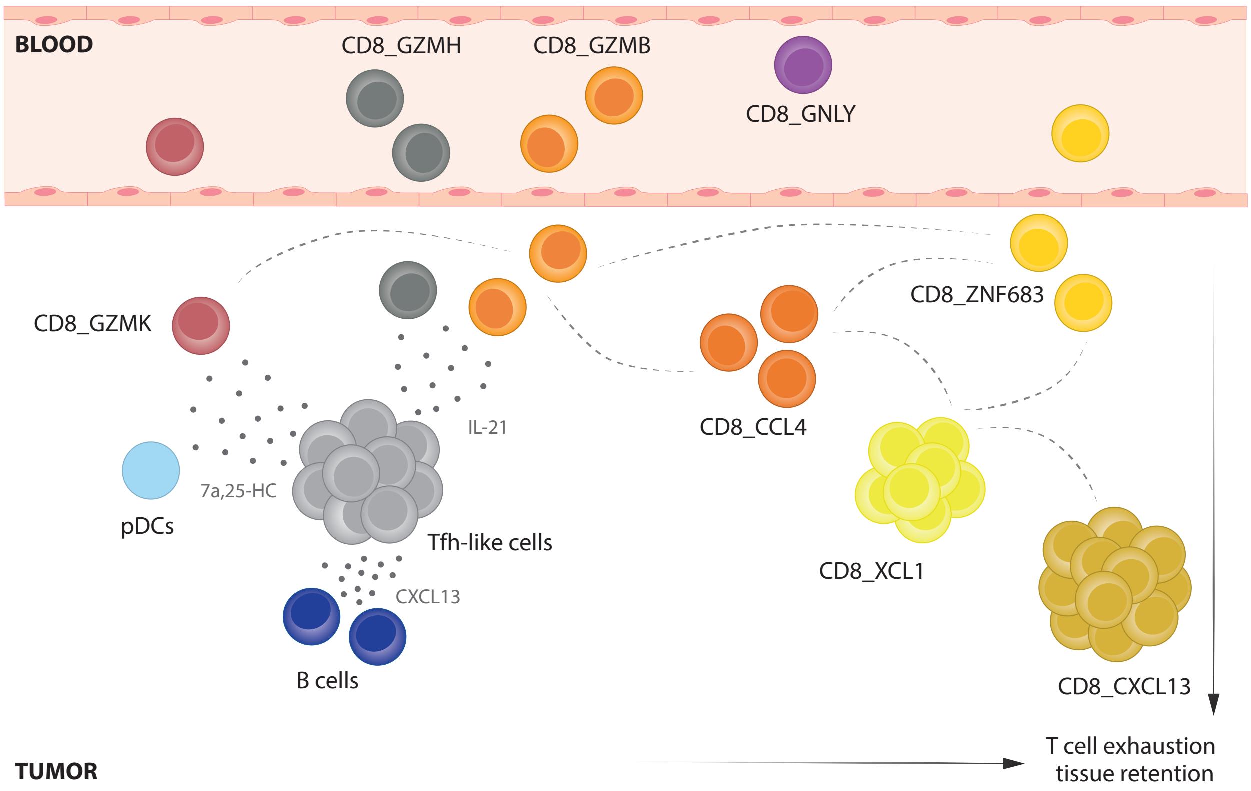


**Fig. S6. Model proposed to explain the lineage relationship between intratumoral and peripheral CD8 T cell subsets in HGSOC.**

Using computational approaches, we inferred the developmental relationship between eight subsets of CD8 T cells and proposed differentiation into exhausted T cells from an intermediate state of effector-like cells. We identified two subsets of recirculating precursor memory-like CD8 T cells, one within a cluster committed to the tissue-resident program (CD8_ZNF683) and another identified as pre-effector cells (CD8_GZMK). Our results suggest the differentiation of cluster CD8_GZMK into the effector-like subset CD8_GZMH. Interestingly, our model proposes a gradual acquisition of the exhaustion program initiated by those effector-like cells (cluster CD8_GZMH) that eventually gives rise to more terminal states with features of tissue residency and chemotaxis (clusters CD8_CCL4, CD8_XCL1, and CD8_CXCL13). Tracking dual-expanded clones and inferring cell-cell interactions, we discovered that *IL21* produced by Tfh-like cells (cluster CD4_CXCL13) is associated with the expression of genes related to the effector function of CD8 T cells. Importantly, Tfh-like cells expressed genes essential for the recruitment of other cells, such as the chemokine CXCL13 and the enzyme CH25H, pointing to their central role in the formation of lymphoid aggregates in ovarian tumors.
